# Population demography maintains biogeographic boundaries

**DOI:** 10.1101/2022.01.13.476105

**Authors:** Chloé Schmidt, Gabriel Muñoz, Lesley T. Lancaster, JP Lessard, Katharine A. Marske, Katie E. Marshall, Colin J. Garroway

**Affiliations:** Department of Biological Sciences, University of Manitoba, Winnipeg, Canada; Faculty of Arts and Sciences, Department of Biology, Concordia University, Montréal, Canada; School of Biological Sciences, University of Aberdeen, Aberdeen, United Kingdom; Department of Biology, University of Oklahoma, Norman, OK, USA; Department of Zoology, University of British Columbia, Canada

**Keywords:** macrogenetics, macroecology, macroevolution, biogeography, population genetics, landscape genetics, conservation, management, mammals, North and South America

## Abstract

Global biodiversity is organized into biogeographic regions that comprise distinct biotas. The contemporary factors maintaining differences in species composition between biogeographic regions are poorly understood. Given the evidence that populations with sufficient genetic variation can adapt to fill new habitats, it is surprising that we do not see more homogenization of species assemblages among regions. Theory suggests that the expansion of populations across biogeographic transition zones could be limited by environmental gradients that affect population demography in ways that could limit adaptive capacity, but this has not been empirically explored. Using three independently curated data sets describing continental patterns of mammalian demography and population genetics, we show that populations closer to biogeographic transition zones have lower effective population sizes and genetic diversity, and are more genetically differentiated. These patterns are consistent with reduced adaptive capacity near biogeographic transition zones. The consistency of these patterns across mammalian species suggests they are stable, predictable, and generalizable in their contribution to long-term limits on expansion and homogenization of biodiversity across biogeographic transition zones. Understanding the contemporary processes acting on populations that maintain differences in the composition of regional biotas is crucial for our basic understanding of the current and future organization of global biodiversity. The importance of contemporary, population-level processes on the maintenance of global biogeographic regions suggests that biogeographic boundaries are susceptible to environmental perturbation associated with human-caused global change.

## Introduction

Since the first voyages of discovery, naturalists and biodiversity scientists have been fascinated by the dramatic faunal and floral transitions we observe among regions (von Humboldt 1807; Sclater 1858; Wallace 1876; Udvardy 1975; Kreft and Jetz 2010; Holt et al. 2013). The factors that differentiate these geographically distinctive species assemblages—*biogeographic regions*—are key to understanding both the current organization of biodiversity and its potential for reorganization under human-caused environmental change. Because biogeographic regions describe general tendencies in the spatial organization of biodiversity, they are essential considerations when prioritizing the conservation and management of global biodiversity (Jenkins et al. 2013). The origins of biodiversity patterns are often viewed as the result of macroevolutionary regional speciation-extinction and colonization dynamics occurring across millions of years (Holt et al. 2013; Lomolino et al. 2016). While these processes underlie the evolution of distinct biotas, they do not explain the processes that sustain regional variation and limit homogenization. The biological constraints that sustain biogeographic boundaries should result from population-level processes that limit species’ abilities to expand into new ecozones. However, the extent to which population-level demographic and genetic processes might scale up to shape continental biotas has yet to be empirically tested.

Biogeographic transition zones are typically characterized by the meeting of distinct biomes or ecozones, and the overlap of various types of habitats that form a patchy environmental mosaic (Ferro and Morrone 2014). We might expect that populations with sufficient genetic variation would be capable of colonizing and adapting to adjacent habitats, eventually causing regional species assemblages to merge. However, we do not generally see such a merger of biotas across biogeographic transition zone habitats. Theory suggests that changes in the demography and genetic diversity of populations associated with such heterogeneous and changing environments could limit the efficiency with which populations adapt to neighboring environments with different conditions (Polechová 2018). Patterns of decreasing population density and effective population size, and increasing genetic differentiation consistently emerge in simulations of population demographics across environmental gradients (Polechová and Barton 2015; Polechová 2018; Bridle et al. 2019). The effective population size is an estimate of the rate at which a population loses genetic diversity due to genetic drift, and it is inversely proportional to the efficiency with which selection can act on beneficial genetic variants (Charlesworth and Charlesworth 2010; Ellegren and Galtier 2016). This theory suggests that range expansion could be restricted because of limits on the efficiency of local adaptation due to the increased strength of drift relative to selection, and the steepness of the environmental gradient (Polechová 2018). Both biotic and abiotic factors contribute to the steepness of environmental gradients (Case and Taper 2000; Goldberg and Lande 2007; Polechová and Barton 2015). In the absence of clines in effective population size, adaptation and spread along environmental gradients remains possible (Kirkpatrick and Barton 1997; Polechová 2018).

We therefore predict that biogeographic transition zones should be characterized by multi-species gradients in the density and genetic characteristics of populations. Assuming underlying environmental gradients are associated with biogeographic transitions, we predicted that effective population size, genetic diversity, and population density would decrease nearer to transition zones, and that genetic differentiation would increase. We tested these predictions for North and South American mammals due to the wealth of demographic, biogeographic, and genetic data available from these regions. Our analyses took advantage of three independently curated open-source genetic and demographic data sets (Lawrence et al. 2018, 2019; Santini et al. 2018a, 2019; Schmidt et al. 2020a, 2020b) and previously described delineations of biogeographic regions (Holt et al. 2013). If our models successfully capture our predicted population-level gradients across these independent data sets, we will have strong empirical evidence supporting the general importance of contemporary population-level processes for preventing the homogenization of communities across biogeographic regions.

## Methods

### Data sources

#### Genetic diversity

We used data from the MacroPopGen database for our estimates of site-level gene diversity (Lawrence et al. 2018, 2019). MacroPopGen aggregates data summaries from the literature for vertebrates in the Americas and includes georeferenced, site-level data for 147 mammal species sampled at 1874 sites across North and South America (Fig. S1). We used the raw site-level estimates of genetic diversity provided on sheet 2 of the Macropopgen database (Lawrence et al. 2018), rather than the re-grouped populations based on genetic differentiation described in their main data set (see next section for reasoning). We selected gene diversity (the average probability that two alleles chosen at random from a sample site are different; Nei 1973) as our metric of genetic diversity because this metric is not strongly influenced by sample size (Charlesworth and Charlesworth 2010), which varies widely in this data set (range: 2 - 1563 individuals per sampling location; mean 48.22 individuals ± 93.51 SD). This and all other population genetic data sets used here are based on microsatellite loci whose diversity are well correlated with genome-wide diversity (correlated at R^2^ ~0.83; Mittell et al. 2015)

#### Effective population size and genetic differentiation

To assess spatial variation in local contemporary effective population size and genetic differentiation, we used a multispecies microsatellite data set compiled by Schmidt et al. (2020a, 2020b) which includes data for 38 mammal species sampled across 801 sites in Canada and the United States (Fig. S1). These data differ from MacroPopGen because they are aggregated raw genotypes instead of compiled literature summaries, which allows users to calculate population genetic metrics that are less routinely presented in the literature. From these data we estimated contemporary effective population size and population-specific F_ST_ (Weir and Goudet 2017).

We estimated the effective population size of the parental generation using the linkage disequilibrium method implemented in the NeEstimator software (Do et al. 2014). Effective population size is reliably measured using linkage disequilibrium (Waples and Do 2010), however, estimates of infinity are returned if populations are very large or if sampling error overwhelms the signal of genetic drift. Sites were excluded from analyses in these cases. We were able to estimate effective population size for 629 sites in 37 species.

We calculated population-specific F_ST_ (Weir and Goudet 2017) using the raw genotypic data in Schmidt et al. (2020a, 2020b). Population-specific F_ST_ estimates the extent of coancestry across all sites in a given species sample—not pairs of sites—and returns a relative, site-level estimate of how far each site has diverged from the common ancestor of populations sampled at all sites. The MacroPopGen data set contains F_ST_ estimates for more populations than the Schmidt et al. data set, but these estimates are summaries of pairwise estimates of F_ST_ for genetic populations defined using a universal threshold that was not suited to our analyses. MacroPopGen F_ST_ estimates are calculated with the extension of pairwise F_ST_ for multiallelic markers like microsatellites (*G_ST_*; Nei 1973), and thus depend on the genetic diversity in the sampled populations. Estimates do not vary between 0 and 1, but have a maximum value of 1-*H_S_* (Charlesworth 1998; Hedrick 1999) where *H_s_* is the mean heterozygosity of subpopulations. This means a universal threshold is incompatible with our analyses because the genetic definition of a population and our interpretation of F_ST_ will vary for each species. For this reason, we use the raw site-level data instead of regrouped populations based on an F_ST_ threshold and recalculate a population-specific F_ST_ metric. Computing population-specific F_ST_ requires at least two sample sites, so we were unable to measure differentiation when the original genotype data were sampled at a single site. Population-specific F_ST_ was estimable for 785 sites in 31 species.

#### Population density

TetraDENSITY (Santini et al. 2018a, 2019) is a global database of >18000 population density estimates (individuals/km^2^) for terrestrial vertebrates. From this data set, we used density estimates for 246 mammal species at 1058 sites in North and South America (Fig. S1). Given the nature of this aggregated data set, species sampled at the same coordinate location sometimes had multiple density estimates for different reasons, including: long-term temporal studies with density estimates across years, multiple methods used to estimate density, or estimates given for multiple localities within sampling areas. These types of studies made up a minority (25%) of the overall data set, and most of the data (88%) had maximum 2 density estimates for species per site. Records for different species collected by different research groups were unevenly temporally sampled, making it impractical to incorporate time into our models. As variation in population density due to temporal change or methodology was not our focus here, we took the average density for species sampled at the same sites. Moreover, sampling method was found to explain little of the variation in population density at broad spatial and taxonomic scales in the TetraDENSITY dataset (Santini et al. 2018b).

We checked the data to ensure there were no island sites where frequent gene flow with continental populations would be unlikely. This was only the case for TetraDensity, and in total we excluded 5 sites that were in the Galapagos, the Caribbean, and Hawaii. We retained the Arctic Archipelago, which is continuous habitat for Arctic species such as polar bears (which were the most consistently sampled species in this region) due to the presence of sea ice.

#### Biogeographic regions

We focused on the biogeographic regions of continental North and South America. We used biogeographic regions previously defined by Holt et al. (2013) to identify transition zones. Holt et al. produced both phylogenetically-based and distribution-based biogeographic regions by clustering mammal species assemblages (i.e. within grid cells). Here, we used the distribution-based maps produced for mammals (see Fig. S6C in Holt et al. 2013), which were generated following procedures similar to those of Kreft and Jetz (2010). We used distribution-based maps because the biogeographic boundaries generated with this approach reflect areas of high overlap in the range limits of species, whereas using the phylogenetic approach, boundaries more likely reflect transition zones at higher taxonomic levels (genus, family, etc.). The distribution-based maps are generated from the clustering of βsim (turnover) values among mammal assemblages, and are robust to changes in data quality and completeness (Holt et al. 2013). In this data set North and South America are comprised of eight biogeographic regions (Fig. S2).

For all sites with estimates of effective population size, genetic diversity, genetic differentiation, or population density, we calculated the geodesic distance (km) to the nearest biogeographic boundary using the dist2Line function in the geosphere package v1.5.0 (Hijmans 2019). We consider biogeographic transition zones and coastlines as biogeographic boundaries. Geodesic distance is an accurate measure of the shortest distance between two points along a curved surface. We computed geodesic distances using the default WGS84 ellipsoid.

### Statistical Analysis

We tested whether distance to the nearest biogeographic boundary was correlated with effective population size, genetic diversity, genetic differentiation, and population density using four Bayesian generalized linear mixed models (GLMM) in the brms package (Bürkner 2019). We ran 4 GLMMs, each with distance to biogeographic boundary as the independent variable, and one of the density or genetic measures as response variables. These data have a hierarchical structure, with sample sites nested within species. We incorporated this structure in our random effect terms: random intercepts for species account for variation in species’ means for each response variable, and random slopes allow the effect of distance to vary across species within the model. Here, species were treated as random samples from a common distribution, so that we can interpret coefficient estimates as the general effect of distance to boundary across all species. If the posterior distribution of the general effect of distance falls entirely above or below zero, this indicates that species have similar positive or negative responses to distance. In contrast, a posterior distribution that overlaps zero may indicate that many species have no detectable response to distance (species-specific coefficient estimates are zero), or that different species have strong positive and negative relationships with distance, generalizing to no overall effect. To visualize results and distinguish between these possibilities, we extracted and plotted species-specific coefficients from the fitted GLMMs. We ran all models with 4 chains with minimum 2000 iterations and weakly informative normal priors on beta parameters with a mean of zero and standard deviation of 1. We used default priors for other model parameters.

Because the nearest biogeographic region boundaries could be either interior region borders or coastlines, we tested whether boundary type affected our results. To this end we classified the nearest boundary for each site as either coastal or interior (Fig. S3). We then re-fit the models described above including a fixed effect for boundary type with an interaction term allowing the effect of distance to vary with boundary type. We included random slope terms for all fixed effects and interactions. Results from models containing boundary type as an interaction term are presented in Table S1.

We tested model residuals for spatial autocorrelation with Moran tests (package spdep; Bivand et al. 2013). The population density model was the only model without significant spatial autocorrelation. We re-ran models for effective population size, genetic diversity, and genetic differentiation using simultaneous autoregressive (SAR) lag models implemented in brms to address spatial autocorrelation. SAR lag models incorporate a spatial weights matrix to account for autocorrelation in the response variable by estimating the strength of spatial dependencies among sites as an additional model parameter.

## Results

Our analyses included gene diversity estimates (Lawrence et al. 2019) from 147 mammal species sampled at 1874 sites across North and South America after filtering, that had a mean of 0.65 ± 0.14 SD (range: 0.04 - 0.94; Fig. 1, Table S2). For population density (Santini et al. 2018a), we included 246 mammal species from 1058 sites (median 9.93; range 0.001 - 11900 individuals/km^2^; Fig. 1, Table S2). We used estimates of effective population size (Schmidt et al. 2020b) for 37 mammal species at 629 sites (median 66.00; 1.00 - 199578.00 individuals per population; Fig. 1, Table S2). Finally, for population differentiation (Schmidt et al. 2020b) we estimated population-specific F_ST_ for 31 species sampled from 785 sites across North America (range: −0.05 - 0.72; mean 0.06 ± 0.08 SD; Fig. 1, Table S2).

**Figure 1.**
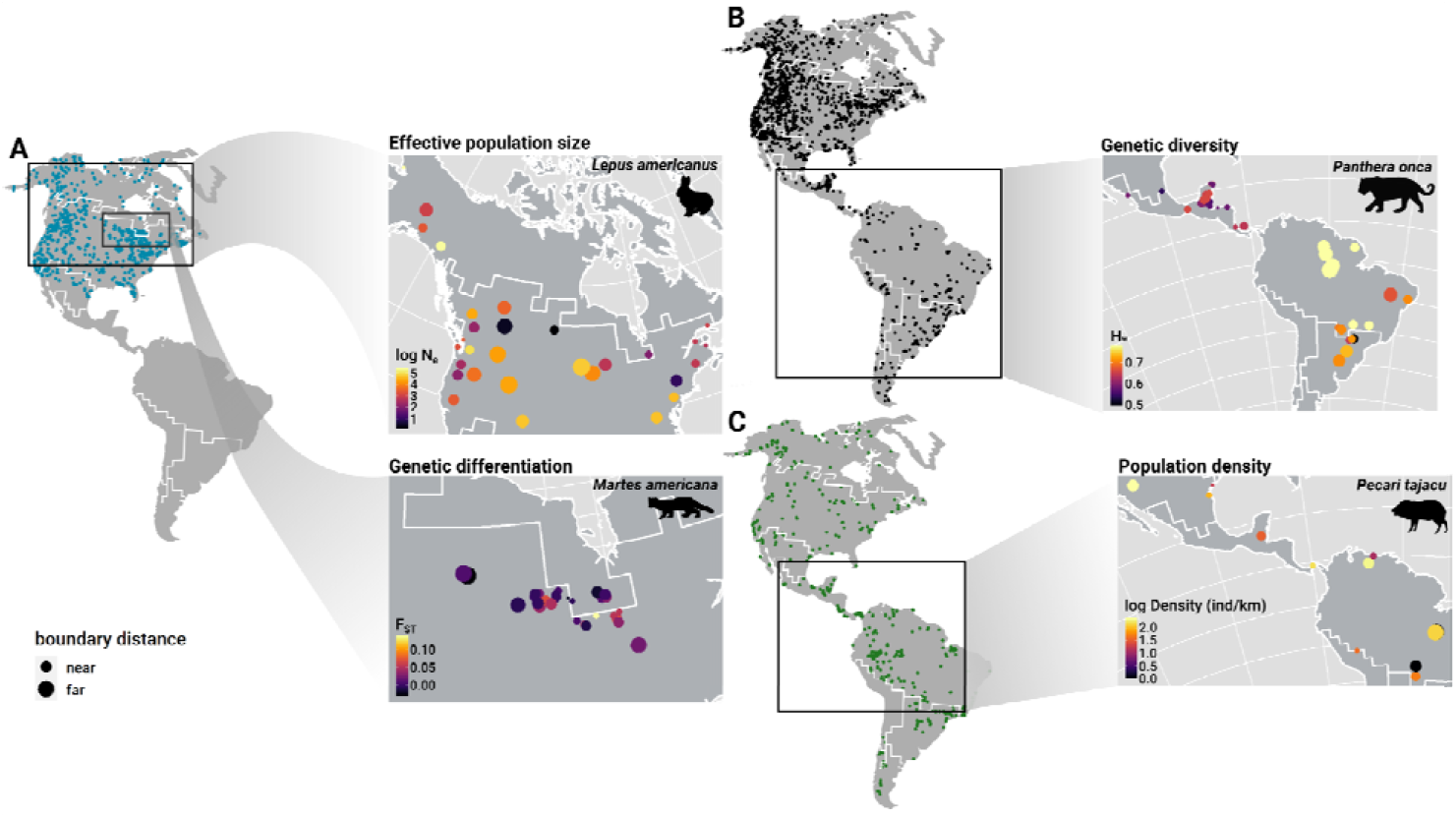
Continental maps show the locations of sites used in this study (A: effective population size (N_e_) and genetic differentiation (F_ST_) estimates from Schmidt et al. data compiled from raw genotypes; B: MacroPopGen genetic diversity (H_e_) estimates; C: TetraDENSITY population density records). One species was sampled at each site. Inset maps show site level values of genetic and demographic variables for select species. The size of points denotes site distance from the nearest biogeographic region boundary.

Genetic diversity, genetic differentiation, effective population size, and population density were all correlated with a sample site’s distance from the nearest biogeographic boundary in our hierarchical regression models (Fig 2; Table 1). In general, as distance from biogeographic boundaries increased, effective population size and genetic diversity also increased, while genetic differentiation and population density decreased (Fig. 2). In other words, genetic differentiation and population density were higher near biogeographic transition zones, while effective population size and genetic diversity were lower. Species by species effects underlying the overall effects (shown in Fig. 2, Figs. S4–S6) trended in the same directions, with no patterns that would suggest moderating effects of species traits or phylogenetic relationships (Fig. S4–S6). Outlier species with strong effects were not consistent across genetic or demographic metrics (Fig. 2). We found no evidence for an interaction between the effect of distance and the type of biogeographic boundary (i.e., whether the nearest boundary was coastal or interior) for genetic variables (Table S1), however there was an interactive effect for population density (estimate = 0.12; 0.01 - 0.23 95% CI; Table S1, Fig. S7). The negative relationship between the nearest distance to transition zone and population density was primarily associated with coastlines, not interior boundaries (Fig. S7).

**Figure 2.**
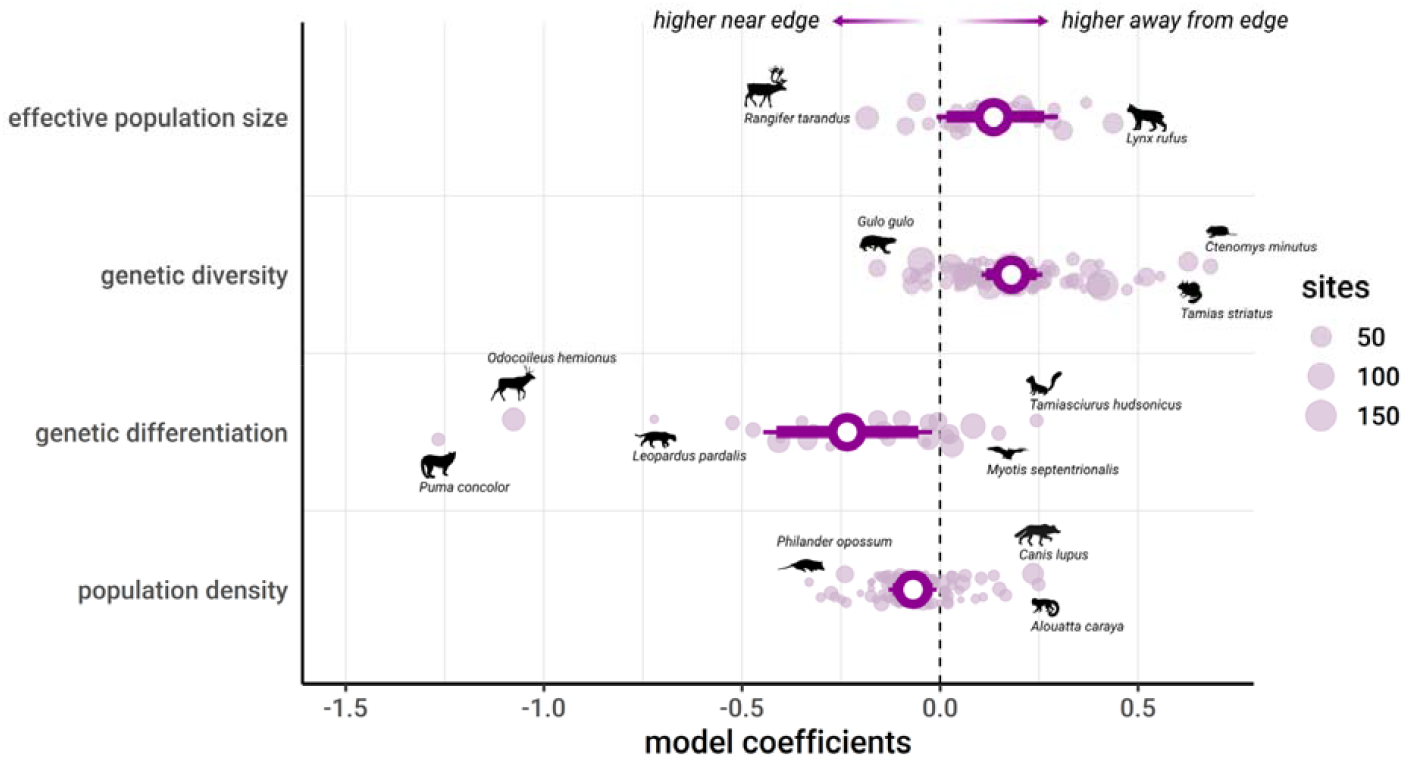
Model coefficients for the effect of distance from biogeographic boundary on population biodiversity variables. Open circles are global coefficient estimates; narrow and thick bars represent 95% and 90% credible intervals respectively. Pale points are the species-specific coefficient estimates that underlie the global estimate, and their diameter denotes the number of sample sites included for that species. Effective population size and genetic diversity increase moving away from region boundaries while genetic differentiation and population density are higher closer to boundaries. Select species at the tails of the distributions of species-specific effects are shown.

**Table 1.**
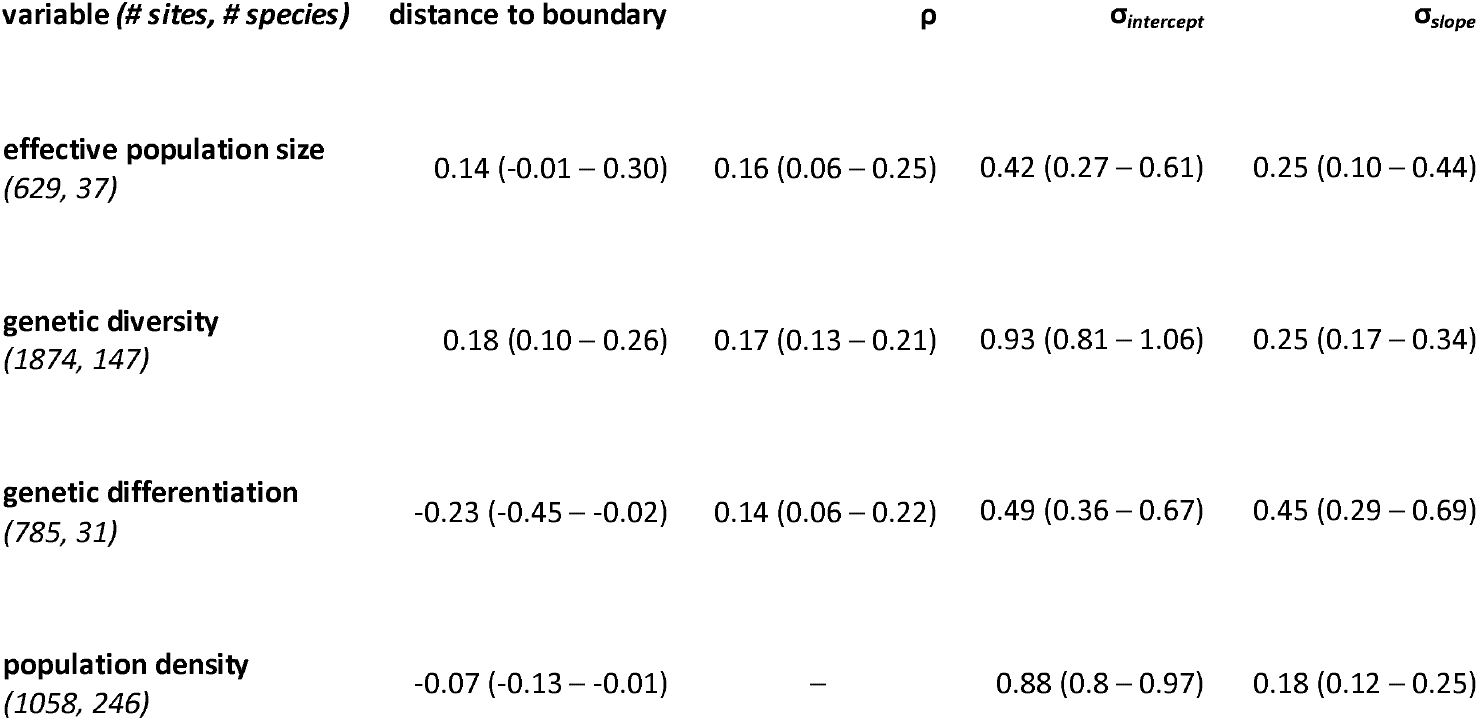
Model summaries for the effect of distance to the nearest biogeographic boundary on genetic and demographic parameters. Effect sizes are given with 95% credible intervals. Rho (ρ) is the coefficient of spatial autocorrelation (for simultaneous autoregressive models only), also presented with 95% credible intervals. Standard deviations (σ) with 95% CIs are given for species random effect intercepts and slopes.

## Discussion

We show that contemporary population demographics, reflected in neutral nuclear genetic diversity and differentiation, vary consistently among species depending on a population’s proximity to biogeographic transition zones. Demography varied in ways that suggest that populations located closer to biogeographic transition zones may be less capable of adapting to the different environmental conditions in and beyond those transition zones (Fig. 2). Stronger genetic drift and reduced adaptive capacity near environmental transitions thus appear to be an important factor in maintaining distinct species assemblages across biogeographic regions.

The spatial organization of global biodiversity results from complex, interacting processes (e.g., historical, evolutionary, ecological) acting over time to shape the biogeographic patterns we observe today. In mammals, biogeographic boundaries are related to tectonic plate movements, and these boundaries are associated with deeper divergences in the phylogenetic relatedness of mammal assemblages across regions (Ficetola et al. 2017, 2021). Climate and elevation have also likely continuously affected dispersal and population demography over long periods to shape regional species assemblages (Ficetola et al. 2021). These historical and contemporary processes have created biogeographic patterns that are, as our results suggest, partly formed by local microevolutionary population processes limiting population spread. Differences in the composition of species assemblages among biogeographic regions thus appear to be maintained by evolutionary limits imposed by increased environmental heterogeneity across hundreds of species at continental extents as predicted by theory (Polechová 2018).

We used globally defined biogeographic regions for mammals and focused on North and South America (Holt et al. 2013). These regions were delineated with a method that maximizes turnover in species composition between regions, and minimizes turnover within regions, such that boundaries represent transition zones where species turnover is rapid. Of course, not all species are restricted to one region. While we identified general genetic and demographic patterns associated with sample location, Holt et al.’s regional delineations did not capture these patterns for all species (Fig. 2). For example, wolverines (*Gulo gulo*) and caribou (*Rangifer tarandus*) had higher genetic diversity and effective population sizes, respectively, nearer transition zones, while American red squirrel (*Tamiasciurus hudsonicus*) populations tended to be more differentiated towards region interiors. These outlier species highlight that there are clearly important species-specific factors that disrupt general patterns in population size and genetic diversity across biogeographic regions (Fig. 2). Future investigations at more local scales (e.g., Morrone 2014) would permit a more focused examination of the environmental or geographic features involved in generating these patterns, albeit with fewer species.

Decreases in genetic diversity across ecological gradients should typically be associated with low population density due to reduced availability of suitable habitat (Polechová and Barton 2015; Polechová 2018). Our empirical findings suggested that population size may be decoupled from density in biogeographic transition zones. Population density tended to increase toward transition zones, but population genetic parameters varied in line with our predictions for decreasing population size (Fig. 2). However, we note that the strength of effect for population density was small and could be spurious. Furthermore, when testing for an effect of boundary type on the relationship between distance and density (Table S1), the interaction term model suggested that this effect was driven by the relationship between population density and distance from coasts (Fig. S7), suggesting that different processes likely not applicable to biogeographic transition zones in general underlie this relationship.

Biogeographic transition zones are often considered conservation priorities because of their high biodiversity (Smith et al. 2001; Spector 2002; Kark et al. 2007). Environmental transition zones and ecotones more generally are sometimes thought of as speciation pumps, where environmental variation and barriers to gene flow create interesting evolutionary arenas with high potential for isolation, differentiation, and speciation (Schilthuizen 2000). In birds, ecotonal zones not only have high species richness by virtue of being areas where different habitats meet, but also harbor high numbers of rare endemic species with narrow ranges (Kark et al. 2007). In light of our results, persistent differences in community composition across biogeographic regions suggest that, if speciation is indeed more frequent in transition zones, these endemic species may be less capable of range expansion due to demographic limits on adaptive evolution. Additionally, one idea in conservation biogeography is that locally adapted populations occupying transition zones may be better equipped to withstand environmental change because they are already adapted to new environments that differ from regional cores (Smith et al. 2001; Spector 2002; Whittaker et al. 2005). From this perspective, biogeographic transition zones would be of high conservation value due to their combination of high species richness, phylogenetic diversity, and populations of genetic significance. However, our findings suggest that prioritizing regional conservation of transition zones over more central locations may run counter to policies intending to maximize genetic diversity and long-term evolutionary potential (Hoban et al. 2020). There will be trade-offs when conserving regions for biodiversity at genetic and species levels. Indeed, spatial correlations between species richness and genetic diversity in general are not straightforward and these two levels of biodiversity tend to be negatively correlated in heterogeneous environments (Schmidt et al. 2022).

Given that contemporary processes are important for the maintenance of diverse species assemblages in biogeographic regions, the persistence of current assemblages may be more susceptible to contemporary environmental change than is currently appreciated. Humans are disrupting environments and the demographics of wildlife populations globally, but whether and how this will affect biogeographic region placement and species compositions is not well known. There is a globally coherent signature of species range movement in response to climate change (e.g., Parmesan 2006). Such range movements can alter the demography and genetics of populations in ways that may lead to the homogenization of both species assemblages and species. For example, climate warming has led to the recent expansion of killer whales (*Orca orca*) into the Arctic where, as a new top predator in that system, they could cause cascading ecosystem changes that might be expected to alter demography and species composition in the region (Lefort et al. 2020). Additionally, most species are evolutionarily young (< 5 million years old) and so may be suspectable to hybridization (Seehausen et al. 2008) when recently diverged species come into secondary contact following range movements related to climate change (e.g., Garroway et al. 2010). This could produce a loss of biodiversity and a merger of gene pools across biogeographic regions. Further, urbanization is now the primary contemporary driver of land conversion for human use (Liu et al. 2020). Urbanization alters the demography, distribution, and genetics of wildlife populations across species in ways that might reshape and reorganize biogeographic regions (Johnson and Munshi-South 2017; Miles et al. 2019; Schmidt et al. 2020b; Schmidt and Garroway 2021). Finally, translocations and invasive species are a major threat to endemics and homogenize biological communities (Capinha et al. 2015; Daru et al. 2021; Yang et al. 2021). Biogeographic region delimitations set our reference points for understanding the distribution and origins of biodiversity and its conservation, thus we need a better understanding of how humans alter their shape and composition.

Through their effects on local population demography, environmental factors appear to set general evolutionary limits across species that contribute to biogeographic patterns at continental scales. Consistent with existing theory (Polechová and Barton 2015; Polechová 2018; Bridle et al. 2019), our results suggest that population demography interacts with environmental transitions in ways that limit population expansion across environmental gradients. This suggests that contemporary microevolutionary processes have contributed to the maintenance of biogeographic regions. Our macrogenetic (Blanchet et al. 2017; Leigh et al. 2021) work adds a bottom-up perspective (i.e., starting at the population-level) to the exploration of biogeographic region formation that has to date been lacking. Population-level microevolutionary processes appear to be fundamental determinants of contemporary biodiversity patterns associated with biogeographic regions.

## Acknowledgements

This manuscript was the result of a working group funded by a Quebec Center for Biodiversity Science grant to JPL and KEM. We thank Laura Pollock, Isaac Eckert, and Federico Riva for comments on the written document and discussion of the topic. We also thank Anna Hargreaves, Brian Leung, Jonathan Belmaker, Lilian Sales, and Shahar Chaikin for additional discussions. We are also grateful to the authors whose work provided the raw data for this synthesis. KEM is supported by a NSERC Discovery Grant. GM and JPL were supported by the Concordia University Research Chair in Biodiversity and Ecosystem Functioning. GM is additionally supported by a Concordia Graduate Fellowship. CS and CJG were supported by a Natural Sciences and Engineering Research Council of Canada Discovery Grant to CJG. CS was also supported by a U. Manitoba Graduate Fellowship, and a U. Manitoba Graduate Enhancement of Tri-council funding grant to CJG. The authors declare no conflict of interest.

## Data accessibility

All data is already in the public domain and sources are cited in the references.

## Supplementary Information

Figures S1-S4

Table S1

**Figure S1.**
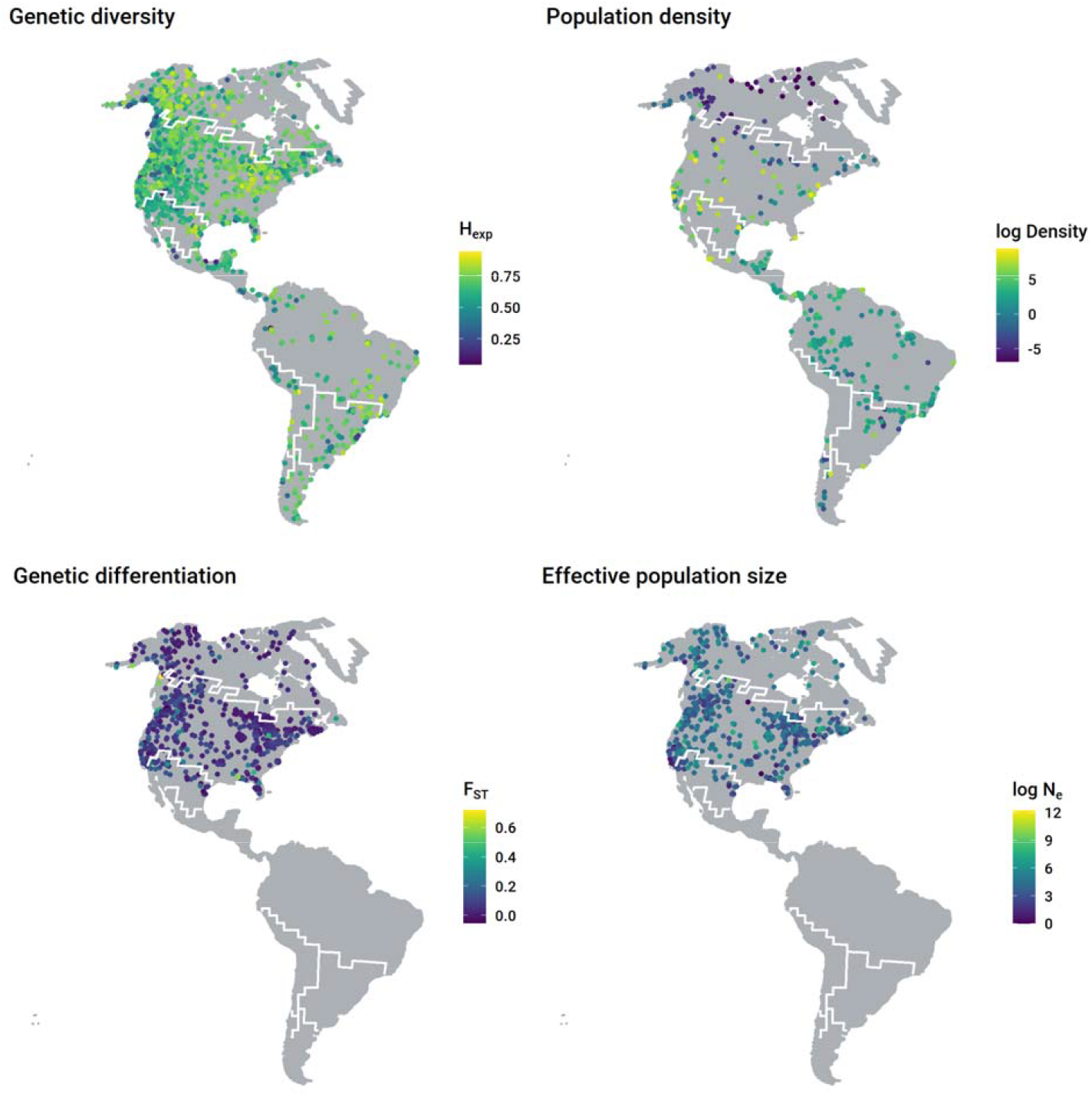
Maps of raw values for population-level biodiversity variables.

**Figure S2.**
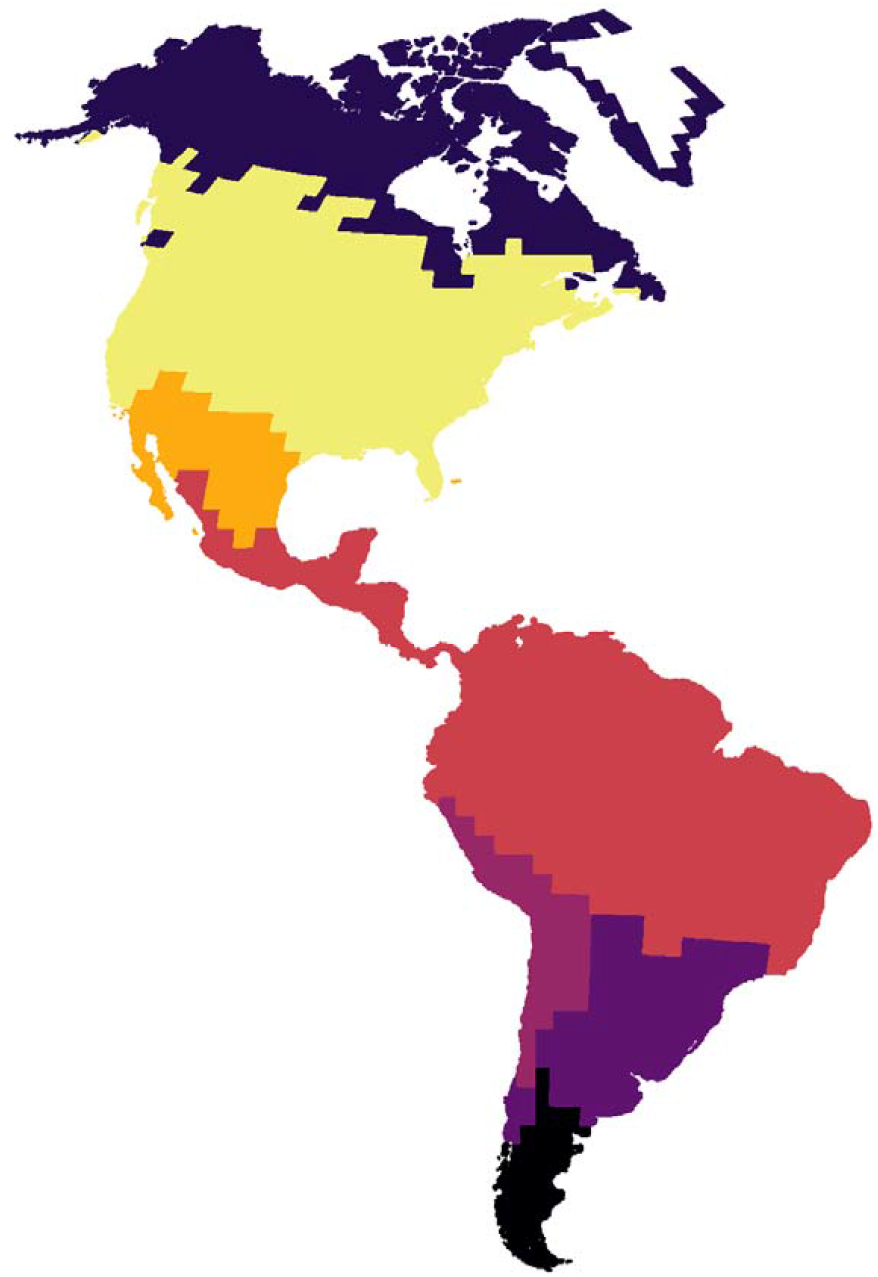
Biogeographic regions of mammals in North and South America (from Holt et al. 2013)

**Figure S3.**
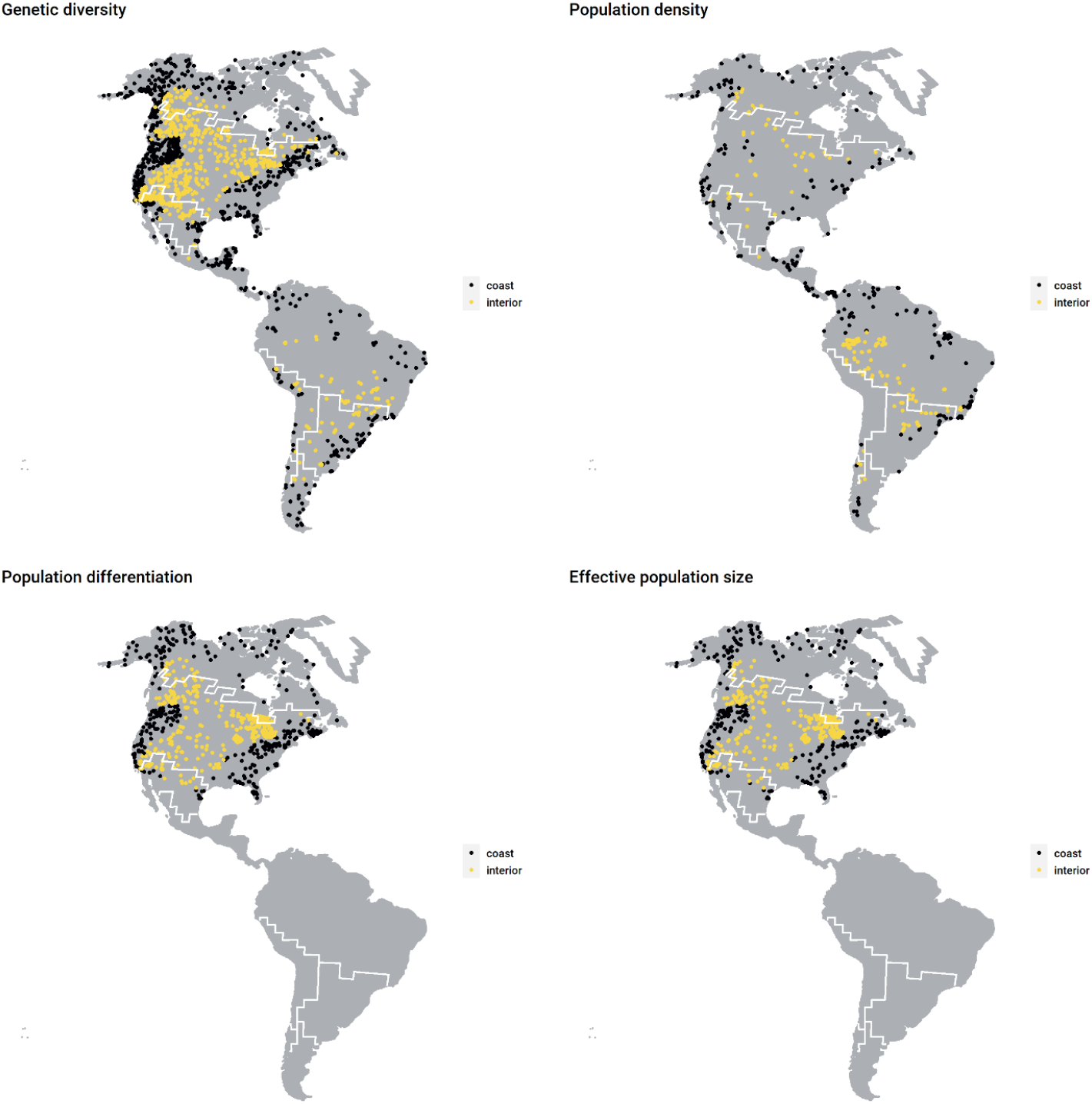
Maps showing whether sample site was nearer to a coastline (black) or interior biogeographic boundary (yellow).

**Figure S4.**
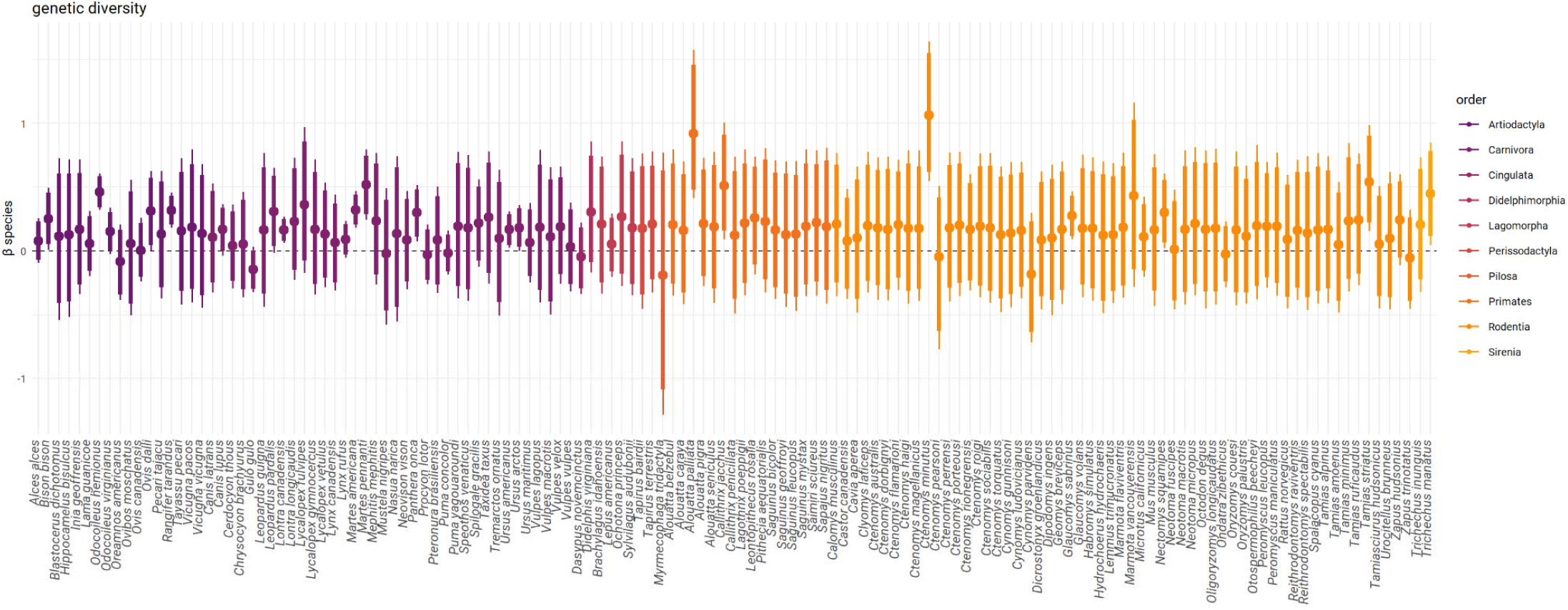
Species-level effects of distance to nearest biogeographic boundary on genetic diversity.

**Figure S5.**
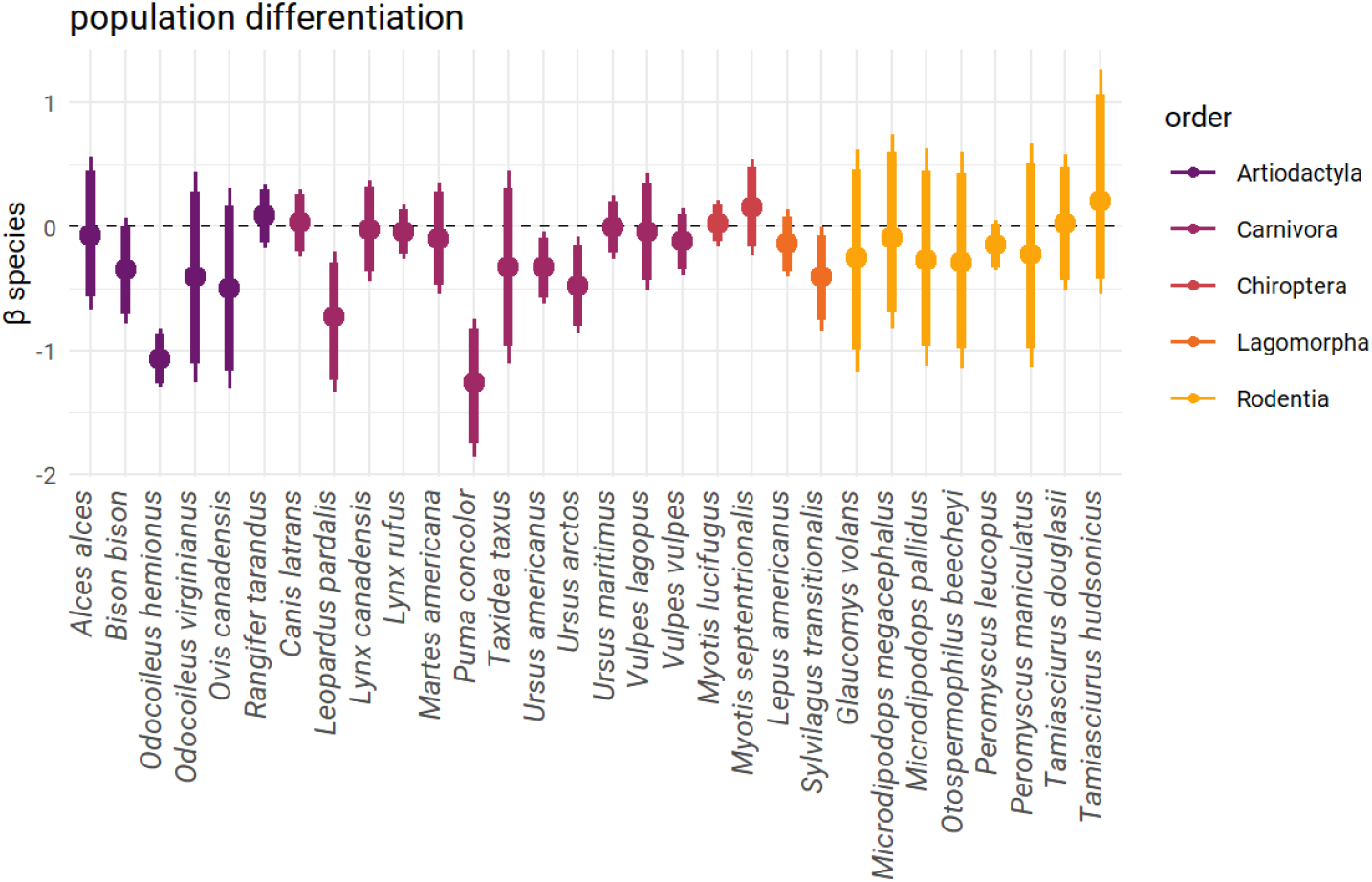
Species-level effects of distance to nearest biogeographic boundary on genetic differentiation.

**Figure S6.**
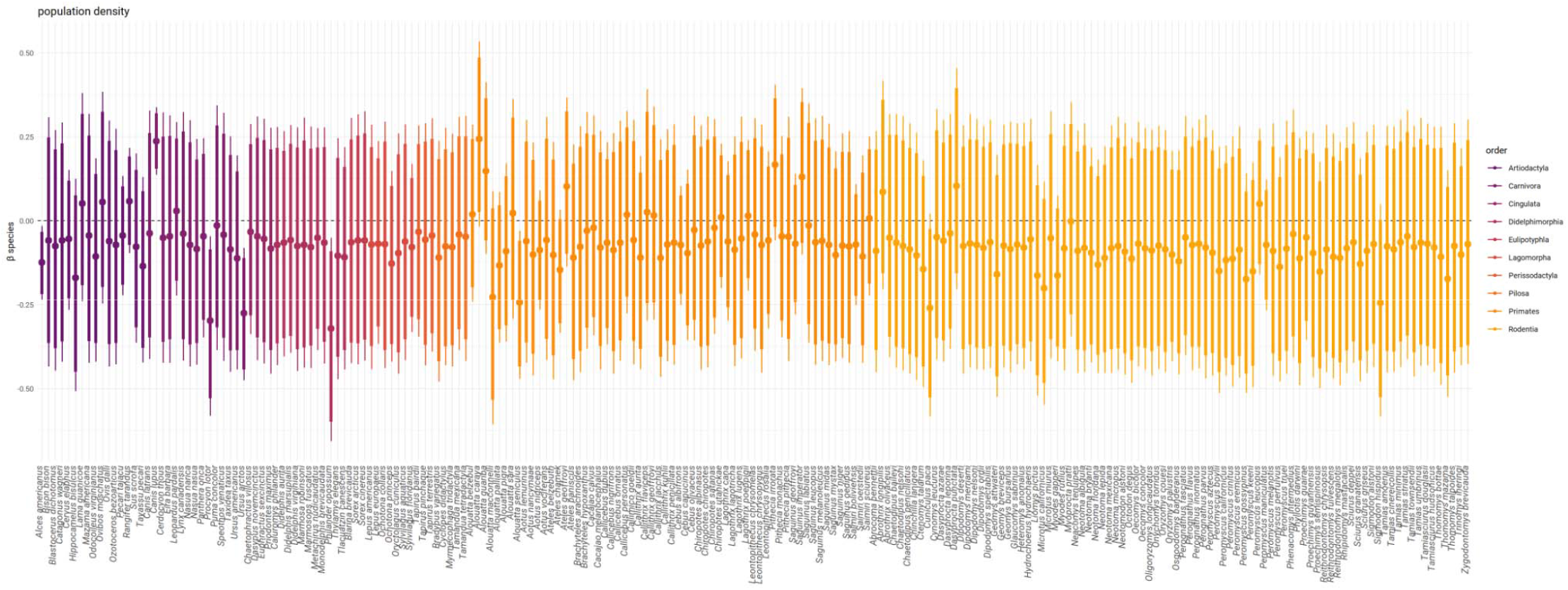
Species-level effects of distance to nearest biogeographic boundary on population density

**Figure S7.**
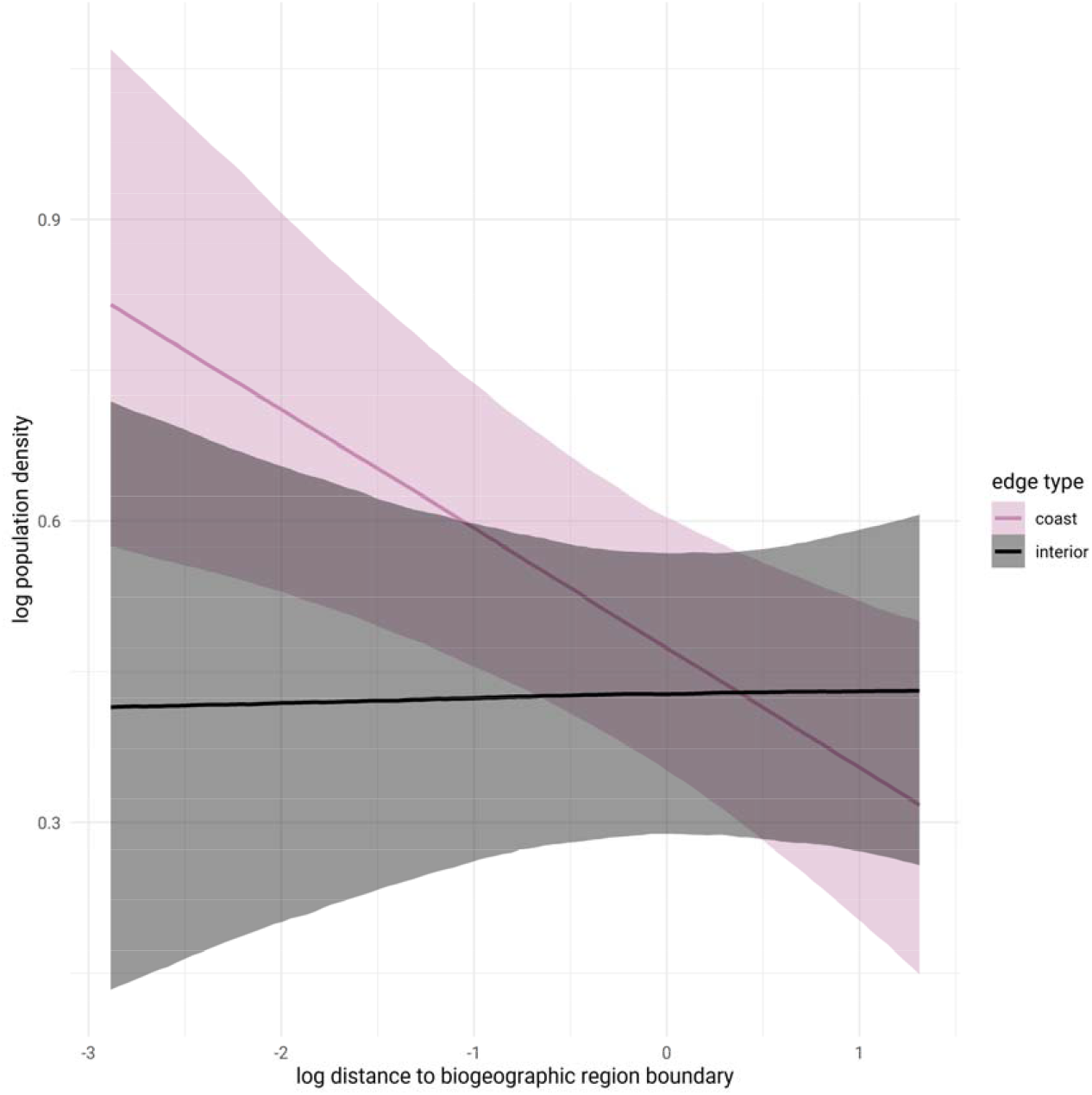
Interaction effect of the type of biogeographic boundary (coastal or interior) on the relationship between the distance to nearest boundary and mammal population density. Increasing population density nearer to boundaries appears to be driven by coastlines.

**Table S1.** Effect sizes and 95% credible intervals of models accounting for boundary type (coastal vs interior). There was no interaction between boundary type and distance (boundary type*distance) for any genetic variable and a weak effect for population density, indicating that the effect of distance only depended on boundary type for population density. The effect of distance on population density was more strongly associated with coastlines (see Fig. S7).

| variable | boundary type * distance | distance to boundary | boundary type: interior |
| --- | --- | --- | --- |
| effective population size | 0.08 (-0.23 – 0.38) | 0.11 (-0.03 – 0.26) | 0.01 (-0.23 – 0.38) |
| genetic diversity | -0.09 (-0.24 – 0.05) | 0.20 (0.10 – 0.29) | 0.10 (0.01 – 0.18) |
| genetic differentiation | 0.32 (-0.11 – 0.71) | -0.27 (-0.60 – 0.06) | -0.17 (-0.46 – 0.10) |
| population density | 0.12 (0.01 – 0.23) | -0.12 (-0.20 – -0.04) | -0.05 (-0.18 – 0.08) |

**Table S2.**
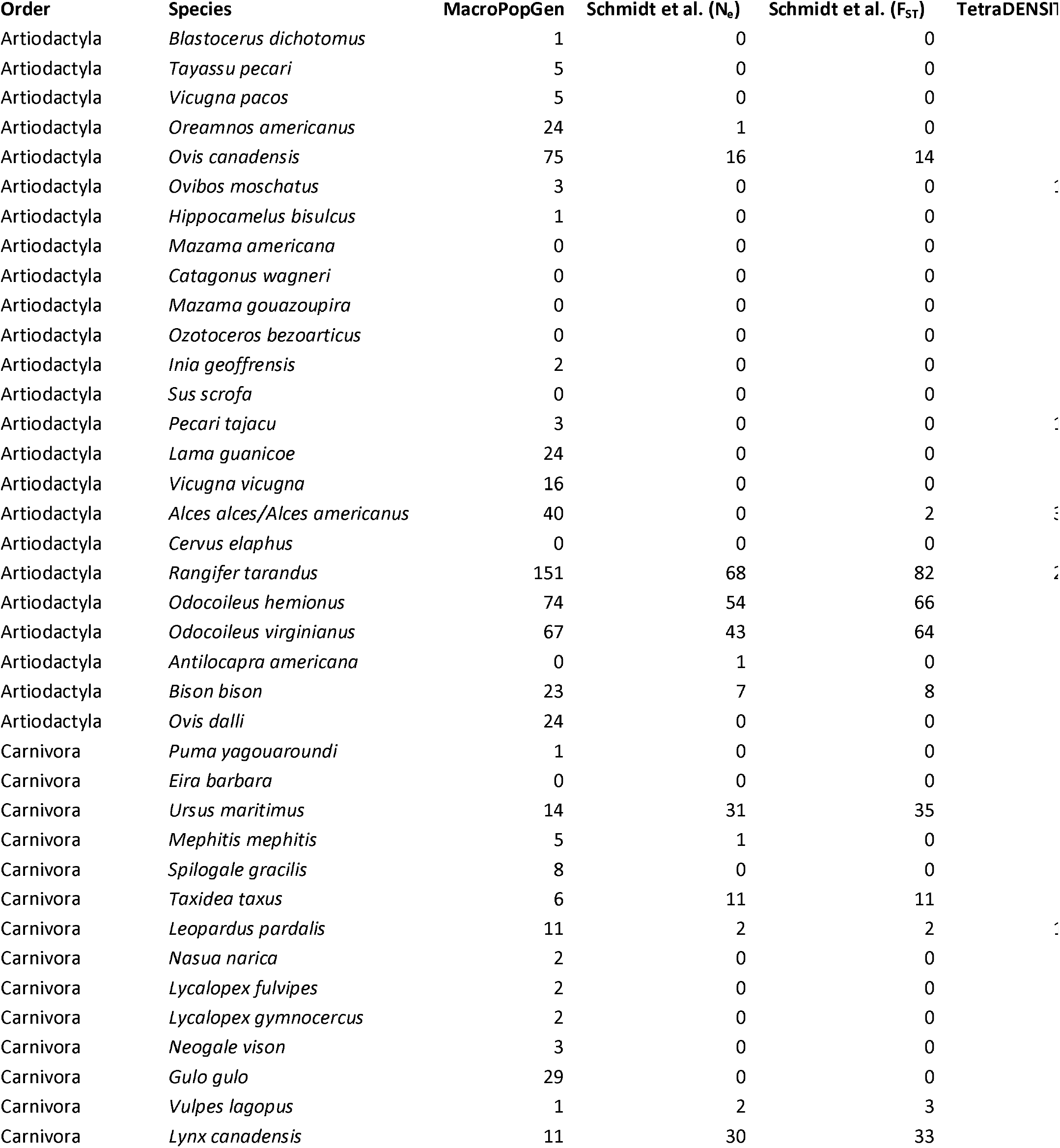
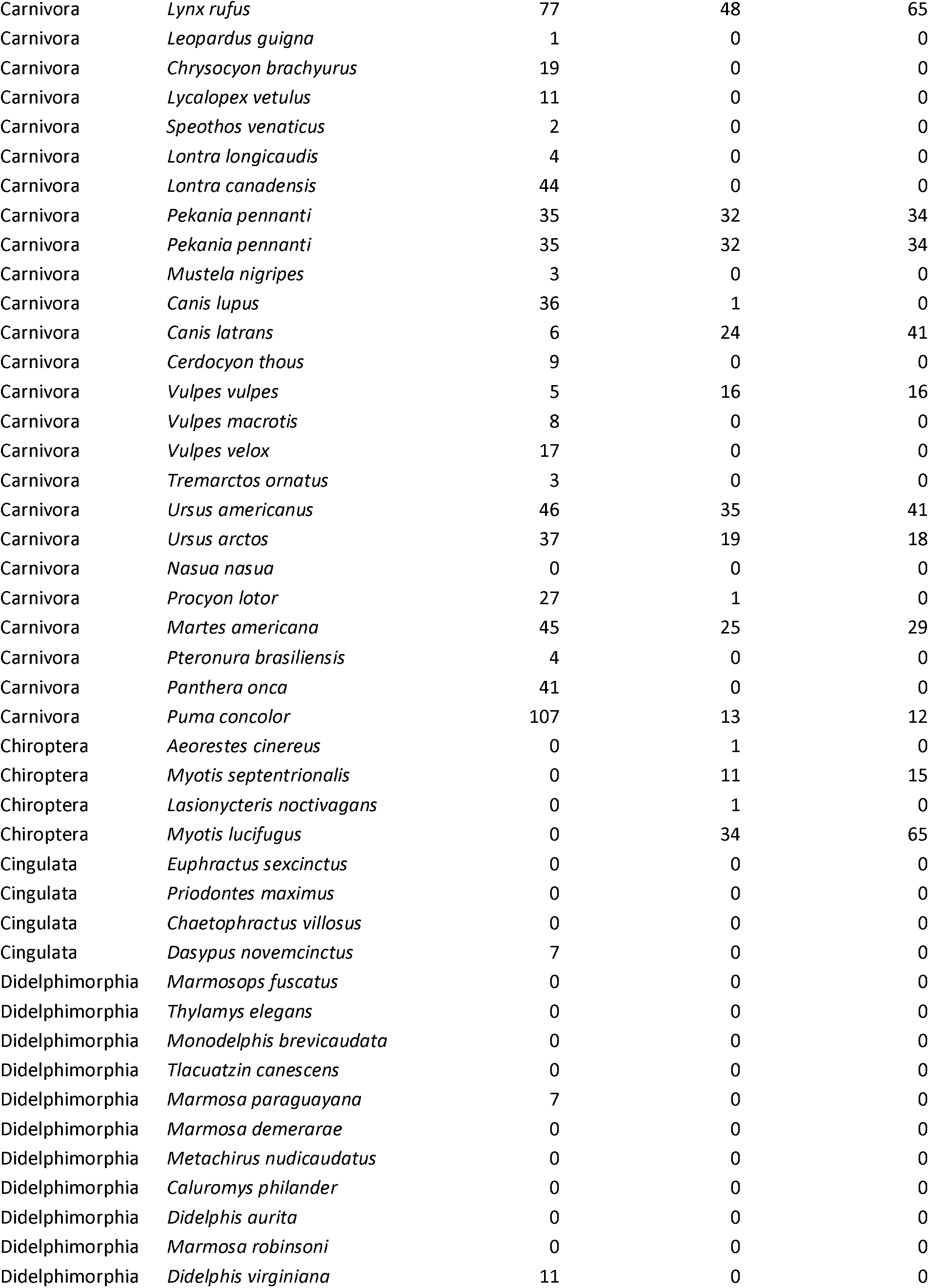
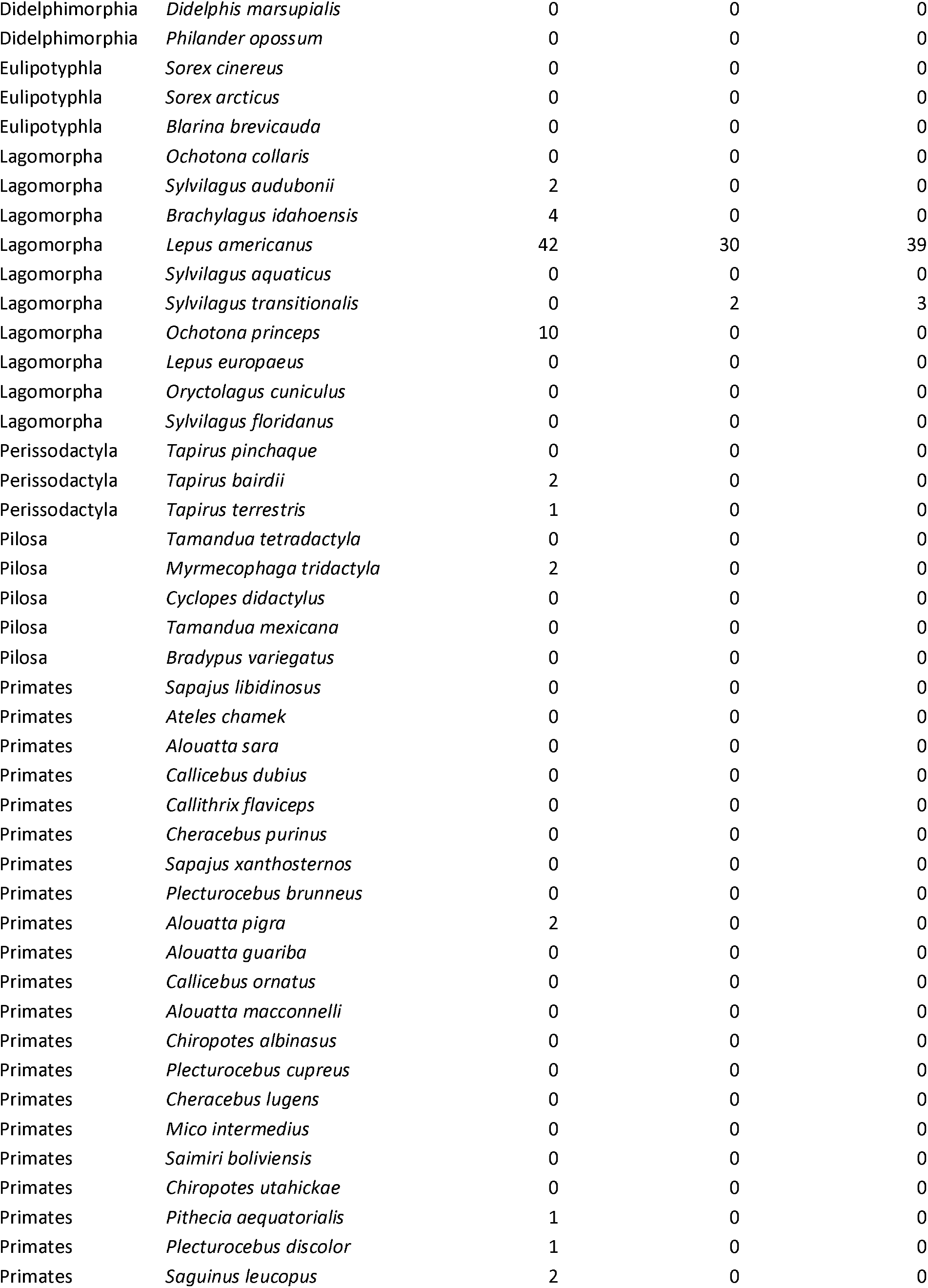
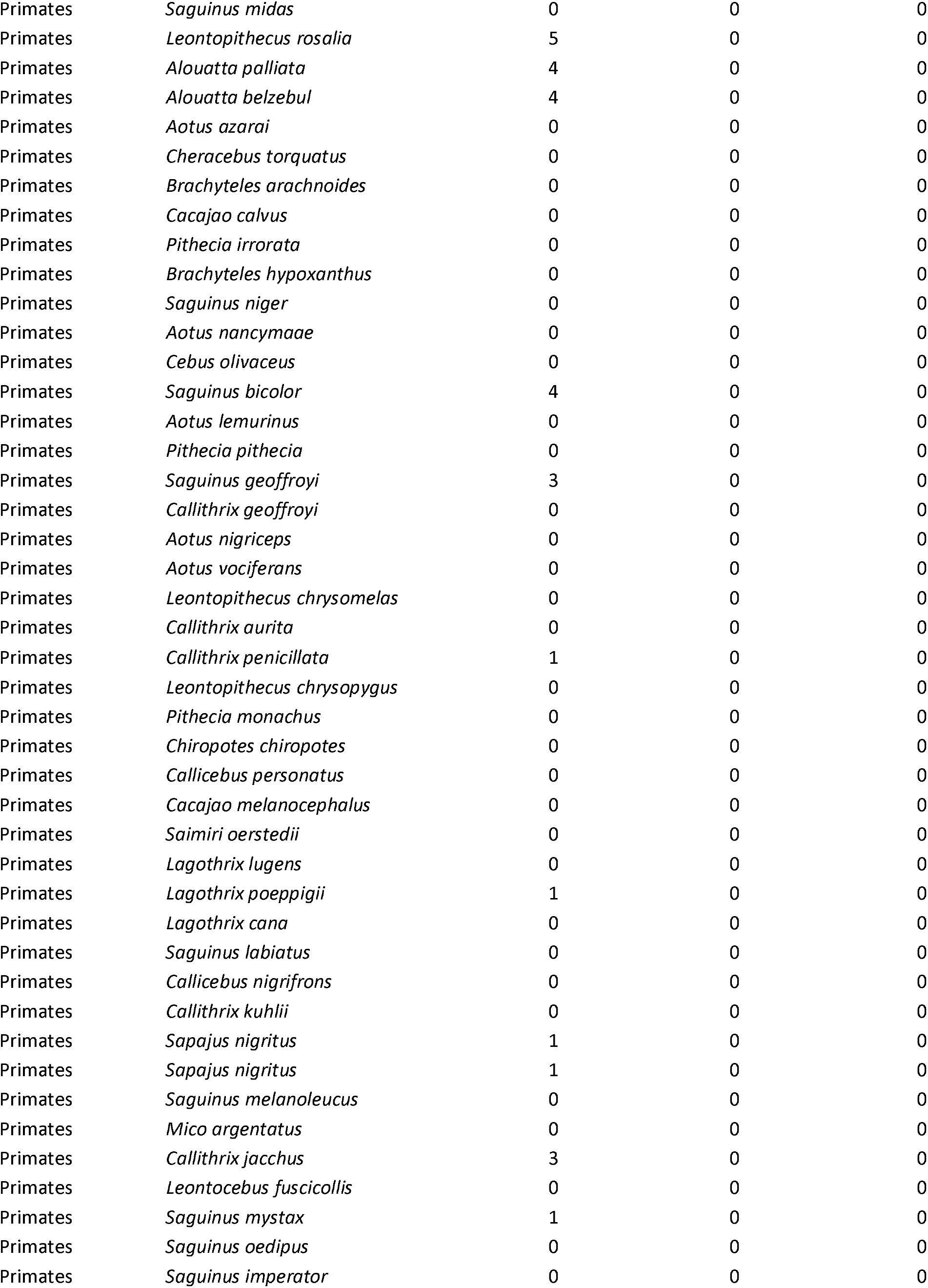
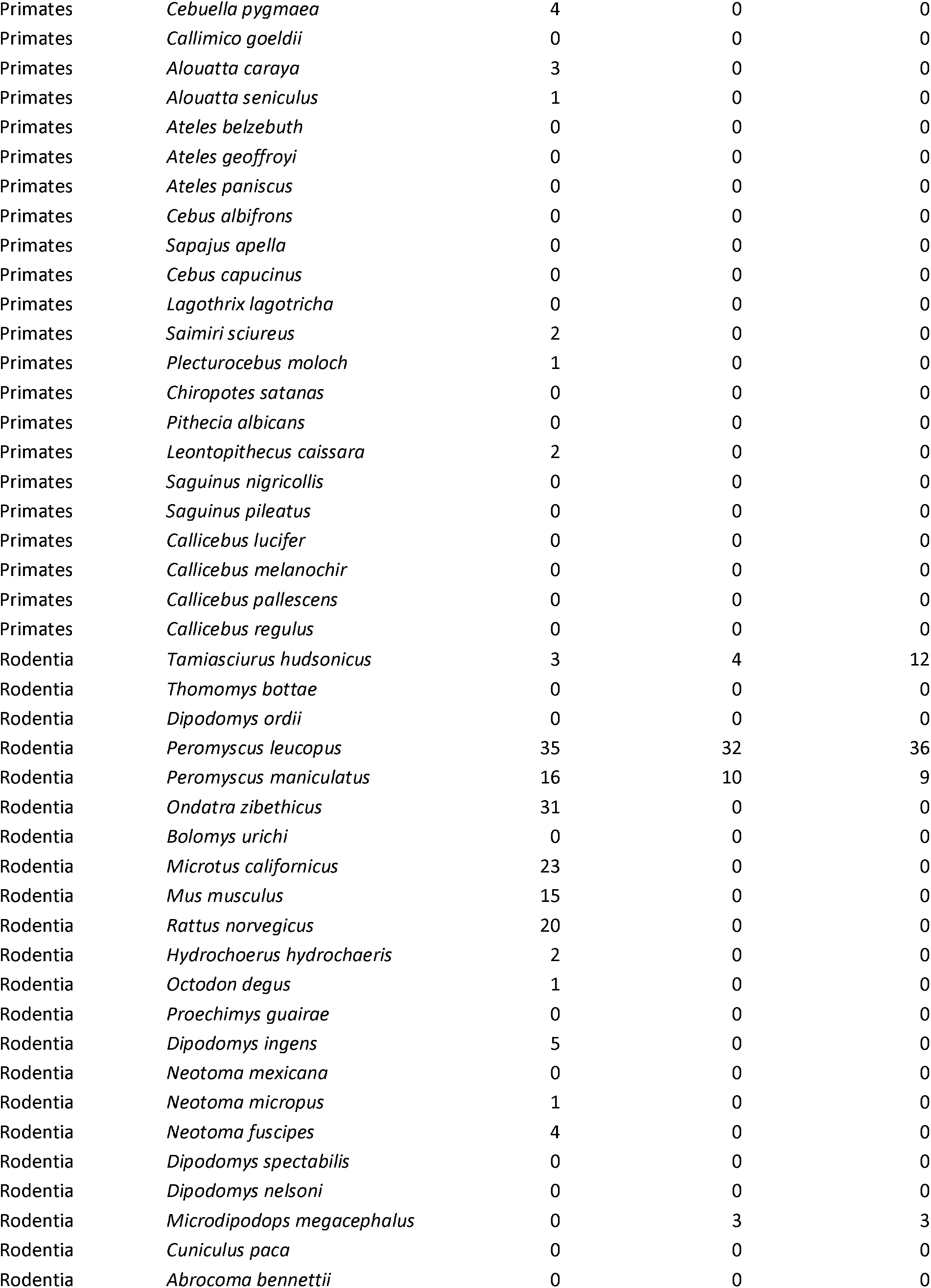
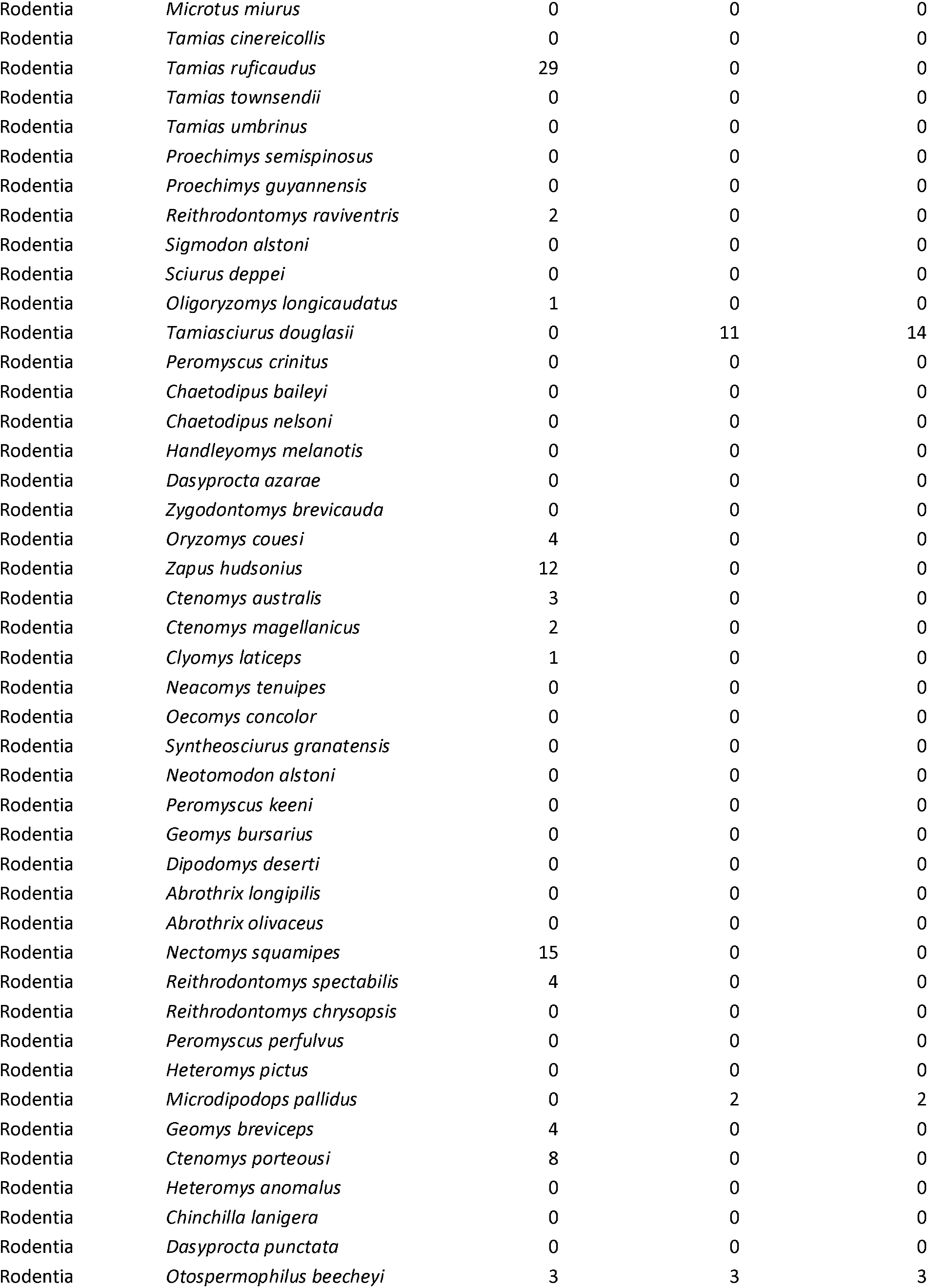
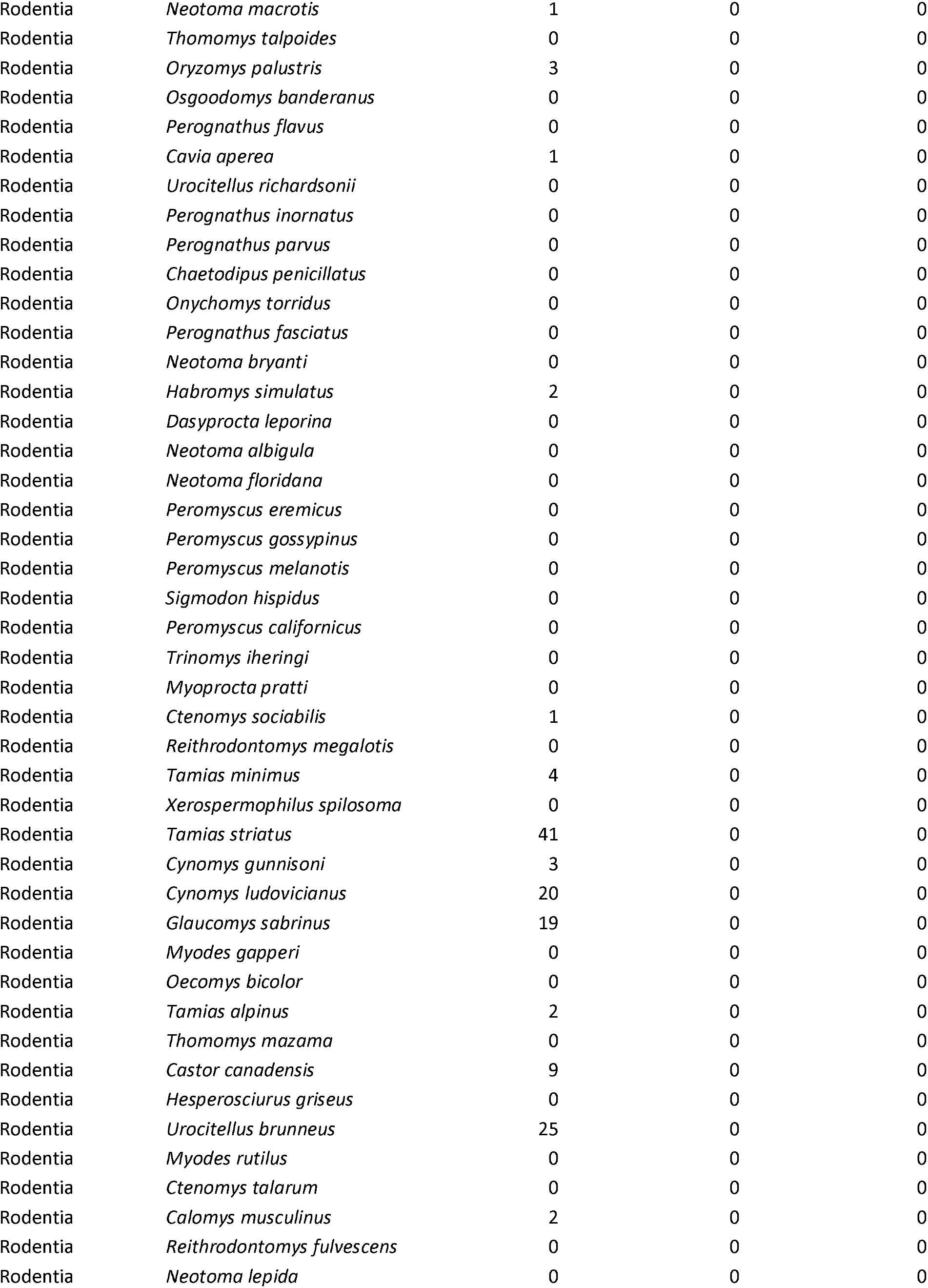
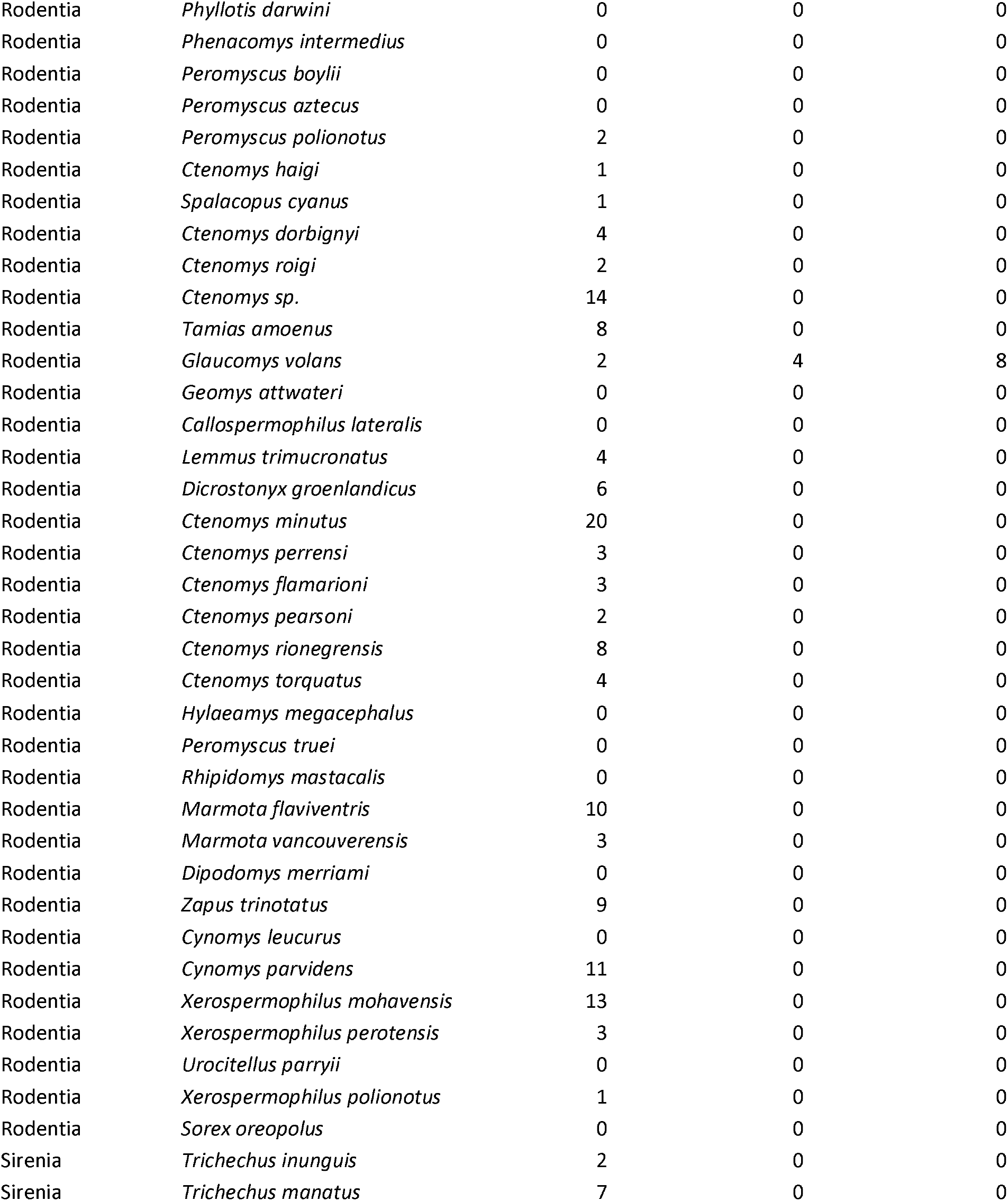
List of species included in analyses and the number of sites per species in each dataset. We used gene diversity estimates from MacroPopGen (Lawrence et al. 2019). Numbers of sites are given separately for effective population size (N_e_) and population differentiation (F_ST_) data from Schmidt et al. (Schmidt et al. 2020). Population density estimates are from the TetraDENSITY database (Santini et al. 2018).

